# The primary transcriptome, small RNAs, and regulation of antimicrobial resistance in *Acinetobacter baumannii*

**DOI:** 10.1101/236794

**Authors:** Carsten Kröger, Keith D. MacKenzie, Ebtihal Y. Al-Shabib, Morgan W. B. Kirzinger, Danae M. Suchan, Tzu-Chiao Chao, Valentyna Akulova, Alesksandra A. Miranda-CasoLuengo, Vivian A. Monzon, Tyrrell Conway, Sathesh K. Sivasankaran, Jay C. D. Hinton, Karsten Hokamp, Andrew D. S. Cameron

## Abstract

We present the first high-resolution determination of transcriptome architecture in the priority pathogen *Acinetobacter baumannii*. Pooled RNA from 16 laboratory conditions was used for differential RNA-seq (dRNA-seq) to identify 3731 transcriptional start sites (TSS) and 110 small RNAs, including the first identification in *A. baumannii* of sRNAs encoded at the 3’ end of coding genes. Most sRNAs were conserved among sequenced *A. baumannii* genomes, but were only weakly conserved or absent in other *Acinetobacter* species. Single nucleotide mapping of TSS enabled prediction of −10 and −35 RNA polymerase binding sites and revealed an unprecedented base preference at position +2 that hints at an unrecognized transcriptional regulatory mechanism. To apply functional genomics to the problem of antimicrobial resistance, we dissected the transcriptional regulation of the drug efflux pump responsible for chloramphenicol resistance, *craA*. The two *craA* promoters were both down-regulated >1000-fold when cells were shifted to nutrient limited medium. This conditional down-regulation of *craA* expression renders cells sensitive to chloramphenicol, a highly effective antibiotic for the treatment of multidrug resistant infections. An online interface that facilitates open data access and visualization is provided as “AcinetoCom” (http://bioinf.gen.tcd.ie/acinetocom/).

## Introduction

In 2017, the World Health Organization (WHO) ranked carbapenem-resistant *A. baumannii* a top priority critical pathogen in the WHO’s first-ever list of priority antimicrobial-resistant pathogens (1). It is estimated that one million *A. baumannii* infections occur worldwide each year, causing 15,000 deaths (2). In the United States, *A. baumannii* is responsible for more than 10% of nosocomial infections, and causes a variety of diseases such as ventilator-associated pneumonia, bacteraemia, skin and soft tissue infections, endocarditis, urinary tract infections and meningitis (3). Despite the urgency to develop new antimicrobial drugs, we know little about *A. baumannii* infection biology and virulence mechanisms because only a few *A. baumannii* virulence genes and their regulation have been functionally studied (4).

Characterizing the transcriptional landscape and simultaneously quantifying gene expression on a genome-wide scale using RNA sequencing (RNA-seq) is a cornerstone of functional genomic efforts to identify the genetic basis of cellular processes (5). Given that RNA-seq identifies transcripts for the entire genome at single-nucleotide resolution in a strand-specific fashion, it is the ideal tool to uncover transcriptome features. Recent technical enhancements to RNA-seq enable improved delineation of transcriptional units to localize transcriptional start sites, transcript ends, sRNAs, antisense transcription and other transcriptome features (5, 6). The ability to quantify gene expression and characterize transcripts from any organism is particularly valuable for the study of emerging bacterial pathogens, where research knowledge lags behind the global spread, health burden, and economic impacts of disease.

To address the lack of basic biological insight of *A. baumannii*, we used a functional genomics approach to investigate the transcriptome architecture of *A. baumannii* ATCC 17978 and to study virulence gene expression and antibiotic resistance in this priority pathogen. *A. baumannii* ATCC 17978 is one of the best-studied strains of *Acinetobacter*, and is a useful model for genetic manipulation due to its natural sensitivity to most antibiotics used in the laboratory (7, 8). Differential RNA-seq (dRNA-seq) facilitates the precise identification of TSS by distinguishing 5’-triphosphorylated (i.e. newly synthesized, “primary” transcripts) and other 5’-termini (such as processed or dephosphorylated) of RNA species (9). dRNA-seq analysis of transcription in virulence-relevant conditions, antimicrobial challenge, and standard laboratory culture generated a high-resolution map of TSS and small RNAs in the *A. baumannii* chromosome, revealing the precise location of 3731 promoters and 110 small RNAs. This comprehensive view of TSS identified two promoters that regulate chloramphenicol resistance by the CraA efflux pump and we discovered growth conditions that stimulate low *craA* expression and render *A. baumannii* sensitive to chloramphenicol. To facilitate similar discoveries by the research community, we created the online resource AcinetoCom (http://bioinf.gen.tcd.ie/acinetocom/) for free and intuitive exploration of the *A. baumannii* transcriptome.

## Materials and Methods

### Bacterial strains and growth conditions

#### Acinetobacter baumannii

ATCC 17978 was obtained from LGC-Standards-ATCC. We used the reference strain containing all three plasmids (pAB1, pAB2, pAB3) for dRNA-seq analysis. RNA-seq revealed that pAB3 was absent in the strain used for early stationary phase transcriptome analysis, indicating that pAB3 was lost during shipping of ATCC 17978 between our laboratories. Comparing RNA-seq reads to the reference genome did not detect any point mutations in the derivative strain lacking pAB3.

Cells were grown overnight in 5 mL Lennox (L-) broth (10 g L^−1^ tryptone, 5 g L^−1^ yeast extract, 5 g L^−1^ NaCl; pH 7.0), sub-cultured the next morning (1:1,000) in 250 ml flasks containing 25 ml L-broth, and grown at 37°C with shaking at 220 RPM in a waterbath. Early stationary phase (ESP) samples were collected when cultures reached an OD_600_ of 2.0 in L-broth. Cell cultures were exposed to several additional growth conditions and environmental shocks to induce gene expression and obtain RNA for the “pool sample”; these conditions are described in the subsequent sentences and summarized in Table 1. In addition to sampling colonies from L-agar plates incubated at 37°C, cells were sampled at select culture densities during growth in L-broth in shaking flasks (OD_600_ values of 0.25, 0.75, 1.8, 2.5 and 2.8). Exponentially-growing cultures (OD_600_ of 0.3) were exposed to different environmental shocks for 10 minutes, including 250 μL of a 1:100 dilution of disinfectant (Distel ‘High Level Medical Surface Disinfectant’, non-fragranced), hydrogen peroxide to a final concentration of 1 mM, ethanol to a final concentration (v/v) of 2%, kanamycin to a final concentration of 50 μg mL^−1^, NaCl to a final concentration of 0.3 M or 2,2’-dipyridyl to a final concentration of 0.2 mM. Cells were also cultured in M9 minimal medium containing 1 mM MgSO_4_, 0.1 mM CaCl_2_ and a carbon source of either 31.2 mM fumarate or 33.3 mM xylose. To mimic growth in environmental conditions, cells were grown at low temperatures (25°C) until an OD_600_ of 0.3. Additional cells were also grown at 25°C to an OD_600_ of 0.3 before cultures were switched to growth at an ambient temperature of 37°C for 10 minutes.

### RNA extraction, ribosomal RNA depletion, DNase digestion and the RNA pool

Total RNA was isolated from *A. baumannii* ATCC 17978 cells using TRIzol as described previously for *Salmonella enterica* serovar Typhimurium (10). The RNA quality was analyzed using an Agilent Bioanalyzer 2100 and RNA concentration was measured on a NanoDrop™ spectrophotometer or the Qubit™ (Invitrogen). The ESP RNA samples were depleted for ribosomal RNA using the RiboZero rRNA Removal Kit Bacteria (Illumina), and contaminating DNA was digested with DNaseI using the TURBO DNA-free™ kit (Ambion). To generate the RNA pool, equal amounts of RNA from each condition were combined into a single sample.

### Library preparation for next generation sequencing

Libraries from three biological replicates of the ESP condition were prepared using the NEBNext™ Ultra RNA Library Prep Kit for Illumina (NEB) according to the manufacturer’s instructions. These libraries were sequenced on an Illumina MiSeq (paired-end reads). The libraries of the pooled RNA sample for dRNA-seq were prepared by vertis Biotechnologie AG (Freising, Germany) and sequenced on an Illumina NextSeq machine (single reads). For dRNA-seq, RNA samples were digested with Terminator Exonuclease (TEX) prior to cDNA library preparation (9, 11). We noted that for highly-expressed genes in the ESP samples, there were distinct transcripts mapped to the opposite strand that perfectly mirrored the expected transcript. This is best explained by incomplete depletion of non-template strand (second strand incorporated dUTP) by the USER enzyme during library preparation with NEBNext™ Ultra. Therefore, we considered only the pooled RNA-seq data from Vertis Biotechnologie AG, which is generated by an RNA adaptor ligation step before cDNA synthesis, to determine the amount of antisense transcription.

### Mapping of sequencing reads, statistics and analysis

The original genome sequence for *A. baumannii* ATCC 17978 identified two plasmids (pAB1 and pAB2) (7). However, it was subsequently revealed that ATCC 17978 carries a third plasmid (pAB3) that was incorrectly assembled as chromosome sequence in the first assembly (12), thus the *A. baumannii* ATCC 17978-mff genome (accession: NZ_CP012004.1) was used as a reference sequence in our study. DNA sequencing reads were mapped using bowtie2 with the–local–very-sensitive settings for use of soft-clipping and maximum sensitivity (13). Conversion into BAM and BigWig files was carried out using SAMtools (14) and Galaxy at galaxy.org (15). Mapped reads were visualized in the Integrated Genome Browser (16) and Jbrowse (17). Number of sequence reads mapped were (the underscore indicates the biological replicate) ESP_1 (10,588,403; mapped uniquely: 8,403,658), ESP_2 (9,374,114; mapped uniquely: 6,769,516), ESP_3 (11,121,847; mapped uniquely: 7,891,371, RNA_pool (66,853,984: mapped uniquely: 19,607,931) and dRNA-seq_Pool (77,237,102; mapped uniquely: 30,041,320). The difference in percentage of uniquely mapped reads of the ESP conditions and the Pool samples is caused by the fact that RNA from the ESP condition was depleted for rRNA, while the RNA samples of the RNA Pool were not. FeatureCounts was used to summarize read counts for each annotated feature (18). These values were transformed into transcripts per million reads (TPM) following the formula provided by Li *et al*. (19); (Table S4).

### Data availability

The raw data and normalized mapped reads are available from the Gene Expression Omnibus (GEO) at accession no: GSE103244.

### Identification and categorization of transcriptional start sites (TSS)

Before the identification of TSS, the coverage files for the RNA Pool and TEX-digested RNA Pool sample were normalized to account for different library sizes. The number of mapped nucleotides for each position was divided by the number of uniquely mapped reads of the smallest library (ESP_2) and then multiplied by the number of reads for the library being normalized. The locations of TSS were predicted with TSSpredator (20) using the default settings with the exception of changing the step height option to 0.2, the 5’ UTR length to 400 and the enrichment factor to 1.5. The locations of the predicted TSS were manually inspected in IGB and Jbrowse. TSS that exist ≤400 nt upstream of the start of a coding region and TSS of non-coding RNA genes were classified as Primary or Secondary TSS; in cases where multiple candidate TSS are upstream, the TSS associated with the strongest expression was designated as the Primary TSS, while all additional TSS were designated Secondary TSS. TSS with intragenic location on the sense strand were classified as Internal TSS and when located on the antisense strand as Antisense TSS. TSS were classified as Orphan if they could not be assigned to any other category. Occasionally, manual curation changed the classification of TSS, e.g. five genes with a 5’UTR >400 nt were classified as Primary or Secondary (Table S2).

### Identification of −10 and −35 sigma factor motifs

DNA sequences upstream of all transcription start sites were extracted in two blocks of nucleotides: between position −3 to −18 (−10 block) and between −16 to −50 (−35 block). Each block was searched with the unbiased motif finding program MEME 4.11.2 (21). Each sequence block was searched with the following settings: one occurrence per sequence with a width of 6–12 bp (−10 motif) or 4–10 bp (−35 motif). Sequence logos were generated by WebLogo (22).

### Identification of candidate sRNA genes and prediction of transcription terminators

The identification of candidate sRNA genes was carried out manually in IGB and Jbrowse. A new candidate sRNA gene was assigned if a short (<500 nt) unannotated transcript was observed in the RNA-seq data. The borders of the transcript were informed by the location of TSS determined by dRNA-seq data and the location of predicted Rho-independent transcription terminators using ARNold with the default setting (23).

### sRNA conservation analysis

The conservation of candidate sRNA genes was investigated using GLSEARCH (24). The genome sequences were obtained from NCBI (*Acinetobacter baumannii* Ab04-mff (NZ_CP012006.1), *Acinetobacter baumannii* AB030 (NZ_CP009257.1), *Acinetobacter baumannii* AB0057 (NC_011586.1), *Acinetobacter baumannii* AB5075-UW (NZ_CP008706.1), *Acinetobacter baumannii* ACICU (NC_010611.1), *Acinetobacter baumannii* strain ATCC 17978 (NZ_CP018664.1), *Acinetobacter baumannii* ATCC 17978-mff (NZ_CP012004.1), *Acinetobacter baumannii* str. AYE (NC_010410.1), *Acinetobacter baumannii* IOMTU 433 (NZ_AP014649.1), *Acinetobacter baumannii* LAC-4 (NZ_CP007712.1), *Acinetobacter baumannii* A1 (NZ_CP010781.1), *Acinetobacter baumannii* USA15 (NZ_CP020595.1), *Acinetobacter baumannii* XH386 (NZ_CP010779.1), *Acinetobacter baumannii* TYTH-1 (NC_018706.1), *Acinetobacter baumannii* XDR-BJ83 (NZ_CP018421.1), *Acinetobacter baylyi* DSM 14961 (NZ_KB849622.1-NZ_KB849630.1), *Acinetobacter equi* 114 (NZ_CP012808.1), *Acinetobacter haemolyticus* CIP 64.3 (NZ_KB849798.1-KB849812.1), *Acinetobacter lwoffii* NCTC 5866 (NZ_KB851224.1-NZ_KB851239.1), *Acinetobacter nosocomialis* 6411 (NZ_CP010368.1), *Acinetobacter pittii* PHEA-2 (NC_016603.1), *Acinetobacter venetianus* VE-C3 (NZ_CM001772.1), *Moraxella catarrhalis* BBH18 (NC_014147.1), *Pseudomonas aeruginosa* PAO1 (NC_002516.2), *Pseudomonas fluorescens* F113 (NC_016830.1), *Pseudomonas putida* KT2440 (NC_002947.4), *Pseudomonas syringae* pv. *syringae* B728a (NC_007005.1). The score shows percentage sequence identity with the reference sequence (*A. baumannii* 17978-mff).

### Northern blotting

Northern blotting was performed using the DIG Northern Starter Kit (Roche) using DIG-labelled riboprobes generated by *in vitro* transcription as described earlier and according to the manufacturer’s instructions (10). Templates for *in vitro* transcriptions using T7 RNA polymerase were generated by PCR using DNA oligonucleotides listed in Table S1. Five μg of total RNA was separated on a 7% urea-polyacrylamide gel in 1x Tris-borate-EDTA (TBE) buffer cooled to 4°C. Band sizes on the Northern blots were compared with Low Range single strand RNA ladder (NEB). After blotting and UV crosslinking, the lane with the ladder was cut off, stained with 0.1% methylene blue solution for 1 min at room temperature, destained in sterile water, and re-attached to the membrane before detection of chemiluminescent bands using an ImageQuant LAS4000 imager (GE Healthcare).

### Chloramphenicol sensitivity and *craA* expression analysis

Cells were cultured overnight in L-broth containing 25 μg mL^−1^ chloramphenicol and the next morning inoculated to OD_600_ 0.05 in 20 mL fresh medium and grown to OD_600_ of 0.3 in a 250-mL flask with shaking at 37 °C. Cells were captured on 0.2 μm analytical test filter funnels (Nalgene), washed with M9 containing 20 mM pyruvate as the carbon source and the filter (retaining the cells) was transferred to M9 plus pyruvate. The cells were washed off the filter and inoculated in fresh 20 mL M9 plus pyruvate medium at an OD_600_ of 0.1 and incubated at 37°C with shaking for 5 days. Sampling for qPCR and chloramphenicol resistance was performed at t = 0, 4h, 24h, 48h, and 96h and transcription was stopped by the addition of phenol:ethanol stop solution (25, 26). For qPCR, total RNA was isolated from cultures using the EZ-10 Spin Column Total RNA Mini-Preps Super Kit (BioBasic) and purity and quality was assessed by 1% agarose gel electrophoresis. For each sample, 5 μg total RNA was DNase treated in a 50 μL reaction using the Turbo DNA-free kit (AMBION), and cDNA templates were synthesized by random priming 0.5 μg RNA in a 20 μL reaction using Verso cDNA synthesis kit (Fisher). PCR reactions were carried out in duplicate with each primer set on an ABI StepOnePlus PCR System (Applied Biosystems) using Perfecta SYBR Fastmix with ROX (Quanta Biosciences). Standard curves were included in every qPCR run; standard curves were generated for each primer set using five serial ten-fold dilutions of *A. baumannii* genomic DNA. Quantitative PCR (qPCR) primers are listed in Table S1. Viable counts were performed by serial dilution and drop-plating on L-agar plates without or with chloramphenicol (10 μg mL^−1^); colony forming units were counted after overnight incubation at 37°C.

## Results and Discussion

### Global identification of *A. baumannii* transcriptional start sites

To generate a comprehensive list of transcriptional start sites for *A. baumannii*, we adopted the technique developed for *Salmonella* Typhimurium in which diverse laboratory conditions were used to elicit transcription activation from as many gene promoters as possible (11). Thus, we selected 16 conditions that reflect the environments experienced by *A. baumannii* during infection, environmental survival, and general stress (Table 1). Specifically, *A. baumannii* was cultured in L-broth and sampled throughout growth at OD_600_ 0.25, 0.75, 1.8, 2.5, 2.8, in L-broth at 25°C, in liquid M9 minimal medium with different sole carbon sources (xylose, fumarate), and grown on an L-agar plate for 16 hours. We also isolated RNA from several “shock” conditions, including disinfectant shock, osmotic shock (addition of sodium chloride), limitation of iron availability (addition of 2,2’-dipyridyl), translational stress (addition of kanamycin), oxidative stress (addition of hydrogen peroxide), temperature shift from ambient to body temperature (25 °C to 37 °C), and the addition of ethanol (which increases virulence in *A. baumannii* infection of *Dictyostelium discoideum* and *Caenorhabditis elegans* (27)). Total RNA was isolated from the 16 conditions, pooled, and used as a single sample.

To enrich for primary transcripts, the pooled sample was digested with Terminator Exonuclease (TEX) (9). Over 20 million mapped sequencing reads were analyzed with TSSpredator software (20) and manually inspected with Integrated Genome Browser (IGB) and Jbrowse (16, 17) to define the genomic location of 3731 TSS (Fig. 1). In addition, three biological replicates of RNA isolated from OD_600_ 2.0 (early stationary phase (ESP)) yielded over 6.7 million mapped reads for each of the ESP biological replicates and the >20 million mapped reads for the RNA pool provided sufficient depth to cover the bacterial transcriptome (28).

**Figure 1.**
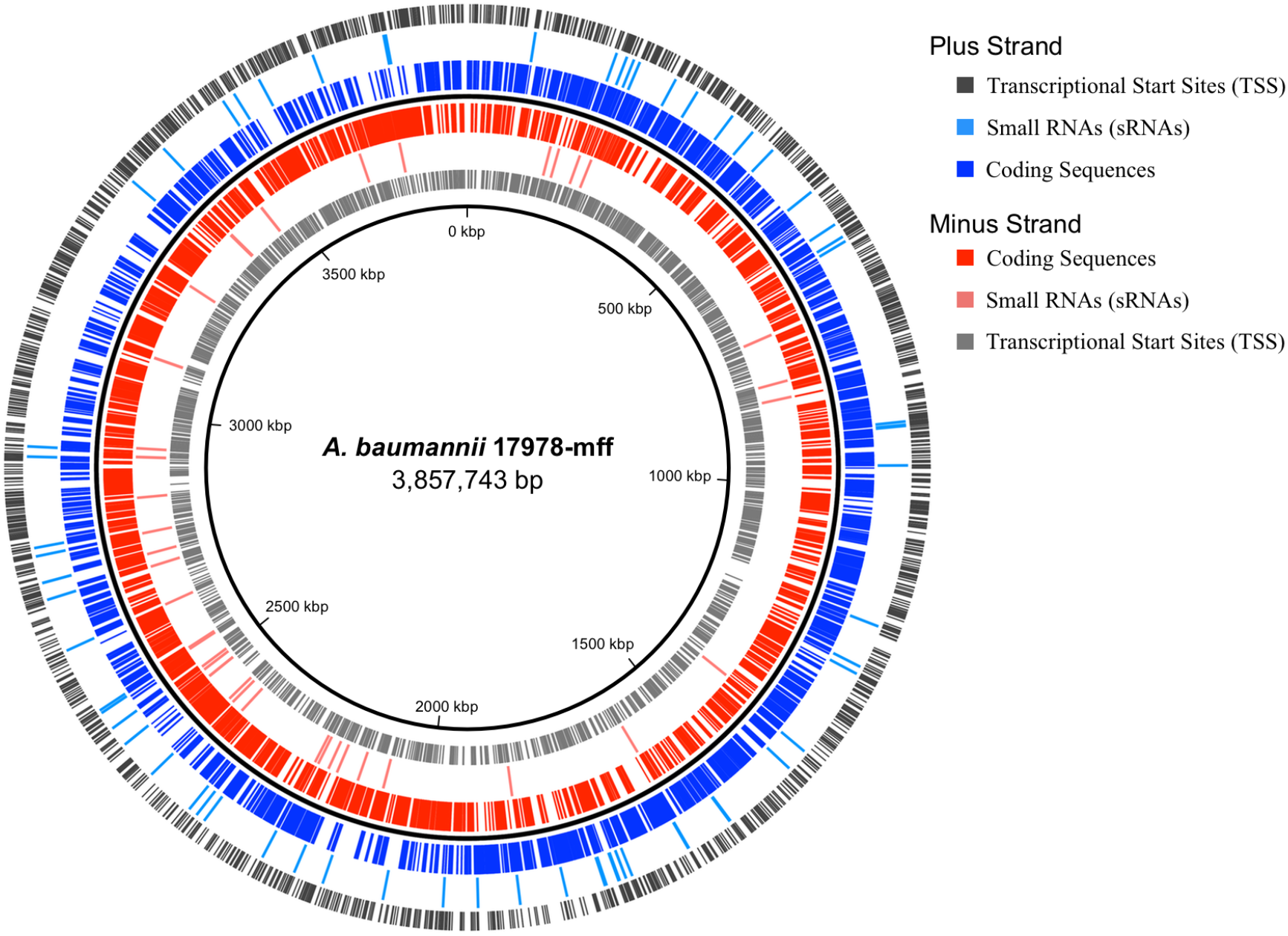
Chromosomal location of coding sequences, small RNAs and transcriptional start sites of *A. baumannii* ATCC 17978-mff. Coding sequences are depicted in blue and red (plus and minus strand), small RNAs in light blue (plus strand) and light red (minus strand) and TSS in grey (outer dark grey ring: TSS on the plus strand, inner light grey ring TSS on the minus strand). The figure was created using Circa (OMGenomics, http://omgenomics.com/circa/).

TSS were categorized according to their locations relative to coding sequences (Fig. 2A). Of 3731 TSS, 1079 (~29%) were primary and 153 (~4%) were secondary TSS (Fig. 2B and Table S2). Manual curation of the TSS predictions determined that five TSS had a 5’ UTR longer than 400 nt; these likely represent unusually long 5’ UTRs, but could encode small open reading frames missed during automated gene annotation (Fig. 2C). About 18% (679) of transcripts were initiated from inside of coding regions (i.e. internal TSS). An internal TSS can drive expression of downstream genes, e.g. the promoter of DNA gyrase B-encoding gene (*gyrB*) lies within the ORF located upstream encoding DNA recombination protein RecF. The biological role of internal promoters that do not drive the expression of a downstream gene is unclear, however, a subset initiates the expression of small RNAs located at the 3’-end of coding genes (see below).

**Figure 2.**
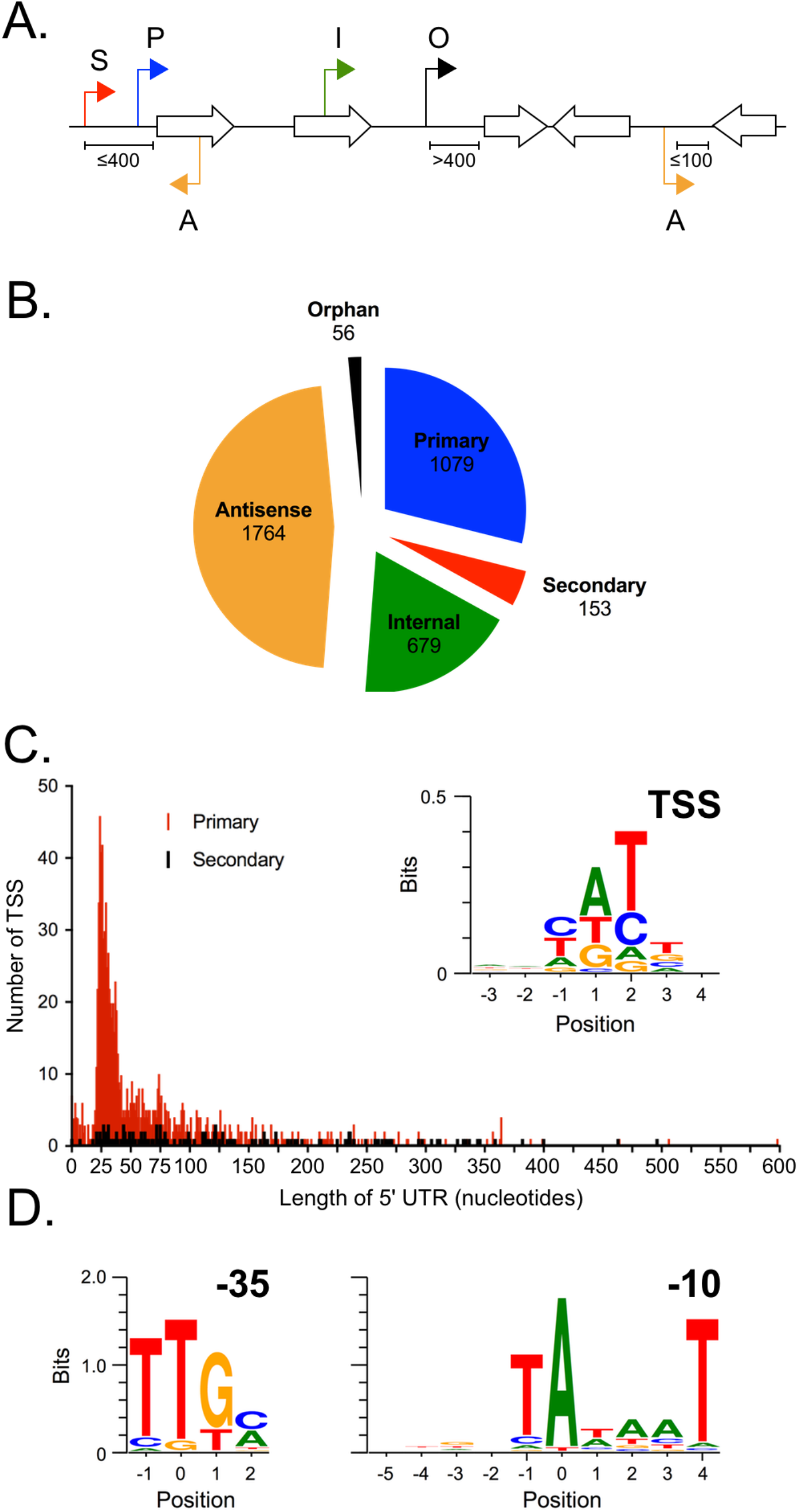
Characterization of transcription start sites. (**A**) Schematic explaining TSS categories. P: Primary TSS, S: Secondary TSS, I: Internal TSS, A: Antisense TSS, O: Orphan TSS. (**B**) Pie chart showing the number of TSS per category. (**C**) Histogram showing the number and length of 5’ UTRs of primary (red) and secondary (black) TSS. The inset illustrates the frequency of occurrence of nucleotides around the transcriptional start site. (**D**) Meme-derived motifs showing −35 and −10 region of *A. baumannii* 17978 promoters.

TSSpredator predicted that almost half (~47%, 1764/3731) of the identified TSS are located antisense of coding regions. Manual inspection revealed that the large majority of antisense TSS produce very short (<100 nt) and low-copy transcripts. The function of these short antisense transcripts is debated and a portion might be considered as non-functional transcripts that are initiated at promoter-like sequences (29, 30). However, an increasing number of studies suggest that pervasive transcription (including short antisense transcripts) exhibit biological roles (31). The ~1.5% (56/3731) of TSS that do not have a clear association with an annotated genomic feature were categorized as orphan TSS.

### *A. baumannii* promoter architecture

To initiate transcription, a sigma factor directs binding of RNA polymerase to DNA within 50 nt upstream of a TSS (32). We determined the consensus promoter structure upstream of the 3731 TSS using unbiased motif searching to detect TSS-proximal and TSS-distal motifs corresponding to the −10 and −35 locations of sigma factor binding sites. Because the spacing between −10 and −35 binding sites differs between promoters, we searched for each motif separately; constraining a specific spacing between −10 and −35 elements prevents resolution of the weaker motifs located around the −35 position. MEME searching identified canonical RpoD (sigma 70) binding site motifs in both the −10 and −35 locations (Fig. 2D), consistent with the expectation that RpoD is responsible for a large majority of transcription initiation in *A. baumannii*. Aligning all 3731 *A. baumannii* TSS revealed that adenine is the most common initiating nucleotide (A=47 %) (Fig. 2C). The majority of transcripts initiate with a purine nucleotide triphosphate (ATP= 47 %, GTP= 23 %); these are the primary energy storage molecules and thus provide a mechanism to regulate transcription initiation. Yet it is position +2 that has the strongest sequence bias (T=57 %). Pyrimidine nucleotides are less abundant in cells, and the preference for pyrimidine nucleotides at the −1 (72 % C+T) and +2 (81 % C+T) positions immediately flanking the TSS may represent a mechanism that reduces accidental transcription initiation from these flanking positions (10).

Untranslated regions (UTR) of mRNA perform multiple biological functions, including providing binding sites for ribosomes and the potential to fold into mRNA secondary structures that regulate transcription and RNA stability (33). The median length of 5’ UTR for primary TSS is 37 nucleotides and secondary TSS is 100 nt (Fig. 2C). Similar UTR analyses have not been performed for the *E. coli* transcriptome; the median length of 5’ UTR in *Salmonella* Typhimurium, another Gamma-proteobacterium, is 55 nt for primary TSS and 124 nt for secondary TSS (10).

### Identification of *A. baumannii* ATCC 17978 small RNAs

Small, regulatory RNAs are a class of post-transcriptional regulators that modulate gene expression of many cellular processes (34, 35). Functions have been determined for some sRNAs in *Enterobacteriaceae* (e.g. *E. coli, Salmonella* Typhimurium), but no sRNA functions have been experimentally determined in *A. baumannii* (4). Two previous studies used bioinformatic and experimental approaches to locate small RNAs in *A. baumannii*. In *A. baumannii* MTCC 1425 (ATCC 15308), 31 sRNAs were predicted bioinformatically, of which three of which were verified by Northern blotting (36). In *A. baumannii* AB5075, RNA-seq analysis of exponential growth in lysogeny broth (LB) identified 78 sRNAs, six of which were verified by Northern blotting (37). Here, we employed dRNA-seq on pooled growth conditions for *de novo* detection of sRNA candidates; pooled RNA samples coupled with the high resolution determination of TSS has been shown to identify a greater number of sRNA per genome compared to traditional methods (11). Using the search criteria described in Material and Methods, sRNAs could occur within intergenic regions, antisense to known coding regions, or within the 3’ end of mRNA. dRNA-seq provided the necessary resolution to detect TSS within the 3’ end of coding genes. We identified a total of 110 candidate sRNAs expressed by *A. baumannii* during ESP growth and in the pooled growth conditions (Table S3). As an example of sRNA annotation, mapped sequence reads of eight sRNAs is depicted in figure 3A. The majority (74/110; 67%) of sRNAs were in intergenic regions. The 3’ regions of coding genes are increasingly recognized as genomic locations that can harbor sRNA genes (5, 38, 39). In *A. baumannii*, we identified 22 sRNAs that mapped to the 3’ regions of coding genes, all of which possess their own promoters located within the upstream coding gene. We anticipate that there will be additional regulatory sRNAs produced by endo-nucleolytic processing of an mRNA, as observed in other bacteria (39). However, additional experiments are required to annotate such sRNAs effectively (39). Only a small number (14/110; 13%) of sRNAs were located antisense of coding genes.

**Figure 3.**
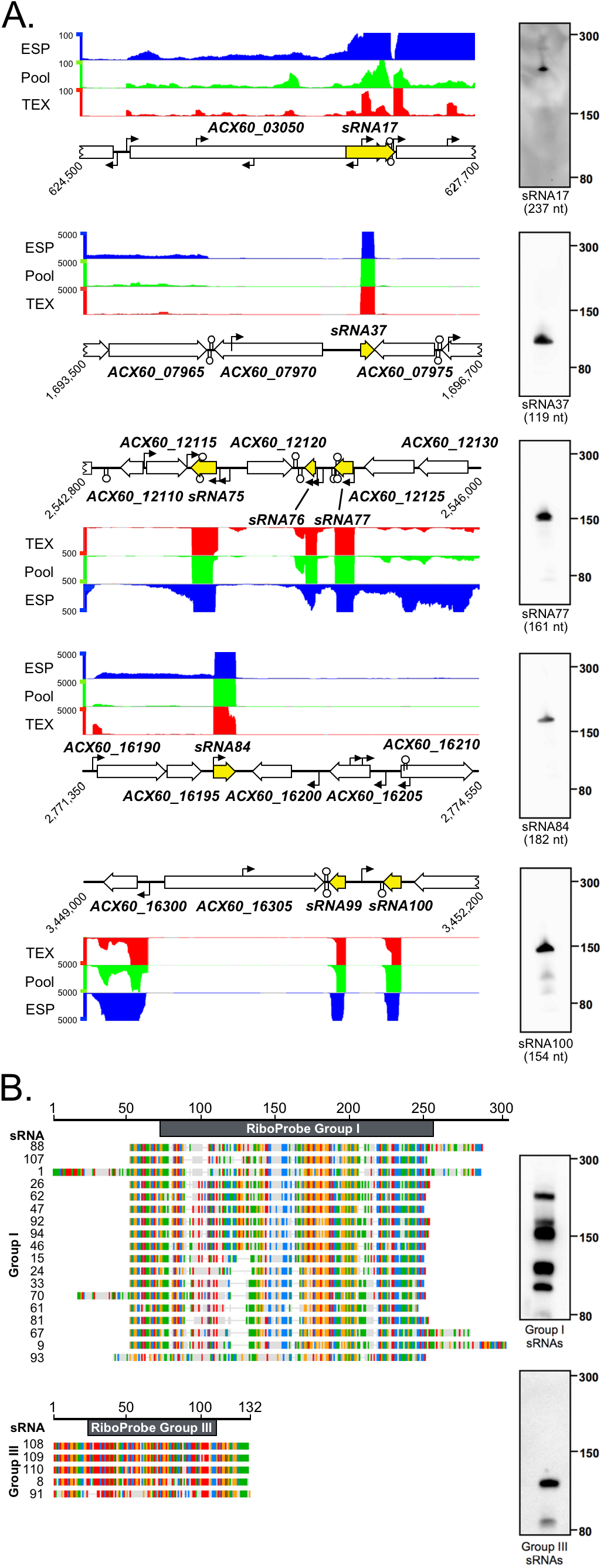
sRNA in *A. baumannii* ATCC 17978. (**A**) Normalized, mapped sequence reads from RNA-seq show the expression of sRNAs 17, 37, 75, 76, 77, 84, 99 and 100 (yellow arrows). Curved arrows depict TSS identified in this study and lollipop structures are predicted rho-independent terminators. Northern blotting of selected sRNAs are shown to the right. RNA was isolated from ESP and five μg of total RNA was loaded per lane. The sRNA sizes below the individual blots have been predicted from (d)RNA-seq data. (**B**) Sequence alignment of Group I and Group III sRNAs created with the Geneious Software (v. 8.1.8); coloured bases indicate conservation in at least 50% of aligned sequences (A, red; C, blue; G, yellow; T, green). The riboprobes used in Northern blotting are depicted as black bars atop the sRNA alignments.

We used Northern blots to validate sRNA annotations (Fig. 3). Sequence-specific riboprobes were designed to hybridize to two families of homologous sRNAs and five randomly selected unique sRNA candidates, including one sRNA (sRNA17) located in the 3’ region of ACX60_03050, a predicted TonB-dependent receptor protein. Eighteen of the 110 sRNA candidates were homologs (i.e. present as 18 copies with highly similar nucleotide sequence), which we classified as “group I sRNAs” (Fig. 3B). Gene duplications are common amongst sRNA species and may benefit the bacterium by allowing for rapid evolution and diversification of gene regulation (34). The transcriptomic data suggested a variable length of the group I sRNAs and Northern blotting with a riboprobe hybridizing to the homologous region of group I sRNAs displayed multiple bands between 100-250 nt in length. The group I family of *A. baumannii* ATCC 17978 sRNAs corresponds to the “C4-similar” group in *A. baumannii* 5075, reported previously to share regions of homology with bacteriophage P1 and P7 (37). Two of the group I *A. baumannii* ATCC 17978 sRNAs contained a 5’ region with low sequence homology; the diversity in this region may provide these particular sRNAs with additional species-specific functions (Fig. 3B).

We next considered orthologs of sRNA that have been characterized in model species like *E. coli*. sRNA37 (119 nt) is predicted to be the 4.5S RNA of the signal recognition particle that is involved in membrane protein targeting. sRNA84 (182 nt) is predicted to be the 6S RNA that can bind and inhibit RNA polymerase in *E. coli* (4).

The RNA binding proteins Hfq, CsrA or ProQ coordinate binding of sRNAs to target mRNAs in *Enterobacteriaceae* (40–42). Of these three sRNA-binding proteins, only Hfq has been studied functionally in *Acinetobacter* spp. and only in *A. baylyi. A. baylyi* Hfq possesses an unusually long, glycine-rich C-terminus that could alter Hfq function compared to what has been observed in model species (43). Thus, post-transcriptional regulation by Hfq and other RNA-binding proteins should be explored in *Acinetobacter* species.

### Conservation of *Acinetobacter* small RNAs

A gene conservation analysis was carried out by comparing the nucleotide sequence identity of the sRNAs identified in 17978-mff with 26 related bacterial genomes. The suite of 27 genomes contained fifteen *A. baumannii* strains, seven *non-baumannii Acinetobacter* species and five more distantly related members of the Pseudomonadales order (Fig. 4). The most highly conserved sRNA across all species were 4.5S RNA (sRNA37), 6S RNA (sRNA84) and to a lesser extent tmRNA (sRNA89). The conservation of *A. baumannii* sRNAs was much higher in the opportunistic pathogens *A. pittii* and *A. nosocomialis* showing a mean sequence identity across all 110 sRNAs of ~87% for *A. pittii* and ~84% for *A. nosocomialis* compared to other *Acinetobacter* species (>70% mean sequence identity across 110 sRNAs). Environmental, non-pathogenic *Acinetobacter* species had lower mean sequence identity. Among the bacterial strains analyzed, eleven sRNAs were highly specific to *A. baumannii* (with less than 70% sequence identity across non-*A. baumannii* strains) and five sRNAs were only present in 17978 (sRNA11, sRNA23, sRNA52, sRNA82 and sRNA83). None of the 110 sRNA were highly conserved (> 75% sequence identity) in the more distantly related *Moraxella* or *Pseudomonas* species. We speculate that sRNAs conserved in *A. baumannii*, but absent from other species, may have evolved for roles in *A. baumannii* virulence or antibiotic resistance that future studies might address. The biological functions of *Acinetobacter* sRNAs may not easily be predicted due to the low level of sequence conservation between *Acinetobacter* sRNAs and well-characterized sRNAs from relatively closely-related *Pseudomonas* species.

**Figure 4.**
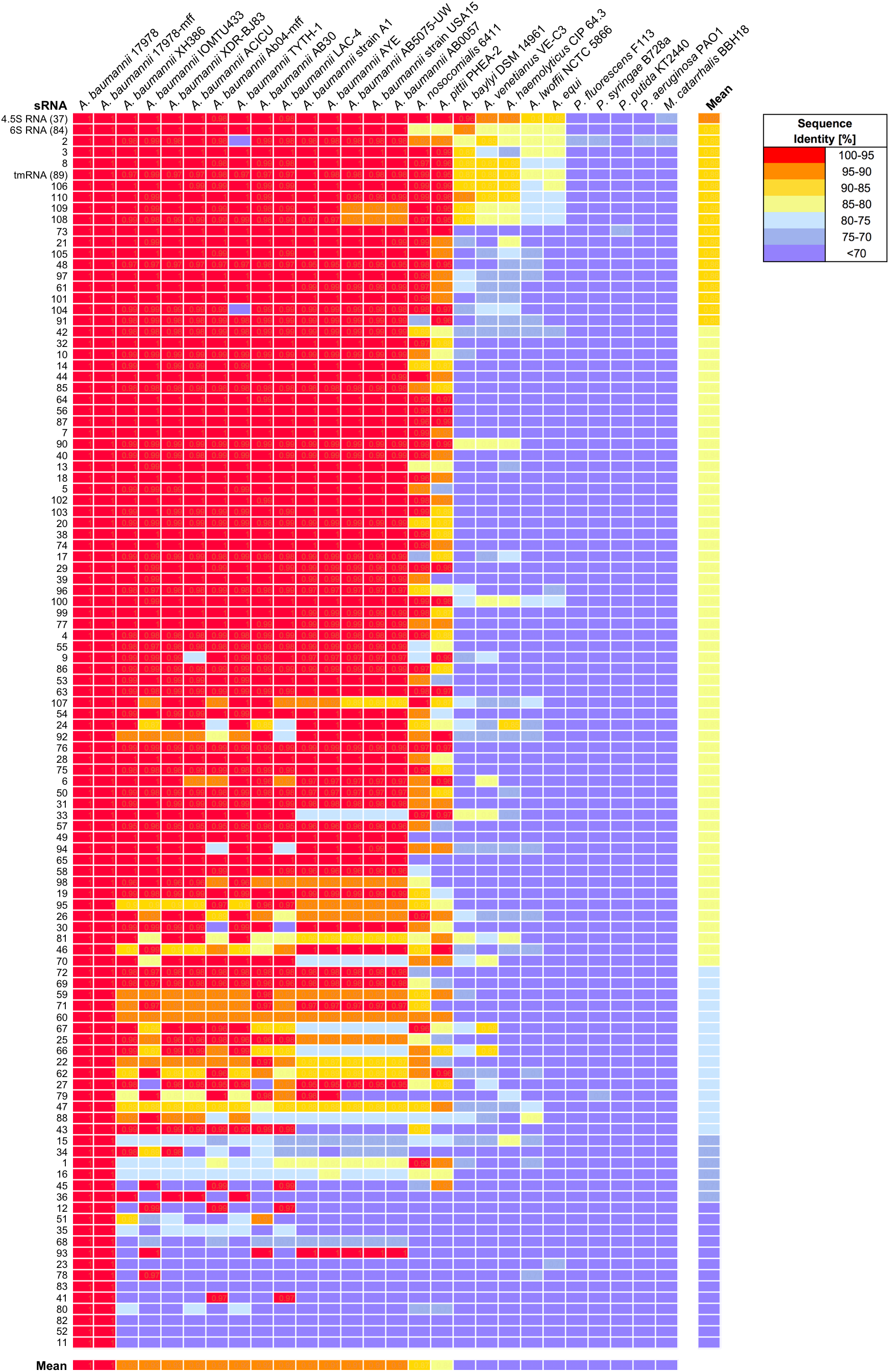
sRNA conservation in representative members of the order Pseudomonadales. Genomes of multiple *Acinetobacter* species, *Pseudomonas* species, and *Moraxella catarrhalis* were compared using GLSEARCH. The colour scale shows the percentage sequence identity of the 110 sRNAs compared to the reference sequence from *A. baumannii* ATCC 17978.

### Functional genomics for the study of virulence and antimicrobial resistance in *A. baumannii*

*A. baumannii* is responsible for an increasing number of hospitalized cases of pneumonia leading to death, which has fueled efforts to identify genes that are important for virulence, host colonization and antimicrobial resistance. In a mouse model of *A. baumannii* pneumonia, genome-wide transposon mutagenesis followed by DNA sequencing (insertion sequencing) identified 157 genes associated with *A. baumannii* persistence in the lung confirming the importance of known virulence genes and identifying several genes with no previous connection to *A. baumannii* infection (44). Our dataset builds upon this analysis by offering detailed molecular insights into the transcriptional architecture of important *A. baumannii* virulence factors. Here, we provide examples of how AcinetoCom can be used by researchers to evaluate transcriptional features of both well-characterized and novel virulence factors, and discuss the importance of *in vitro* growth conditions in the study of virulence gene expression (Table S4).

### Regulation of Zinc Acquisition and the Znu ABC Transporter

Metal acquisition genes are a critical component in *A. baumannii* virulence as they are required to wrest essential co-factors from host chelators (4, 45, 46). Zinc limitation has been demonstrated as a central feature in controlling bacterial replication in the lung and dissemination to systemic sites in the murine model of *A. baumannii* pneumonia (45, 47). In *A. baumannii*, the Zur (<u>z</u>inc <u>u</u>ptake regulator) transcriptional regulator is responsible for mediating the expression of several zinc-associated functions, including the repression of an ABC family zinc transporter encoded by *znuABC*. In *A. baumannii* 17978, the *zur* gene is co-transcribed as part of an operon with *znuCB*, which encode the cognate ATPase and permease protein of the ABC transporter (45). The third component, *znuA*, is found on the opposite DNA strand and encodes a substrate-binding protein (45). During growth in zinc-replete conditions, the Zur protein binds to a 19-bp operator site located 30 nucleotides upstream from the *znuA* translation start site and represses the expression of both *znuA* and the *zurznuCB* operon (45, 47).

TSSpredator identified a total of 8 TSS within the chromosomal region that includes both the *znuA* gene and *zurznuCB* operon (Fig. 5A). Two of the TSS were located 29 and 93 nucleotides upstream of the *znuA* ORF. While both TSS were annotated as antisense to the *zur* gene, their location within the intergenic region also make them prime candidates as primary or secondary TSS to *znuA*. No nearby TSS were predicted for the *zurznuCB* operon; however, manual inspection revealed potential primary and secondary TSS located 21 and 78 nucleotides upstream that were observable but not enriched in the TEX sample relative to the RNApool sample. Based on these predictions, the TSS for *znuA* or *zurznuCB* are found on either side of the Zur-binding site and may represent an important regulatory feature for Zur-based transcription repression (45, 47). An additional TSS was identified within the *znuC* ORF, located 239 nucleotides upstream of *znuB*. To our knowledge, this TSS has not been previously identified in the well-characterized *zurznuCB* operon and suggests the potential for transcription of *znuB* independently from other members within the operon. Such a format of expression may be a mechanism for cells to accommodate modest increases in the intracellular level of zinc while avoiding simultaneous increases in the Zur protein. Further, there are no reports of a Zur box associated with the region surrounding this internal TSS. A Rho-independent transcriptional terminator is also present in this region, located within the *znuB* ORF or 113 nucleotides downstream of *znuC*. Several additional internal TSS were also identified within the *znuA* gene, but did not demonstrate a clear association with downstream genes or with an sRNA transcript. The remaining TSS within the region were annotated as antisense TSS and contribute to the overall theme of pervasive low-level antisense transcription within the *A. baumannii* transcriptome.

**Figure 5.**
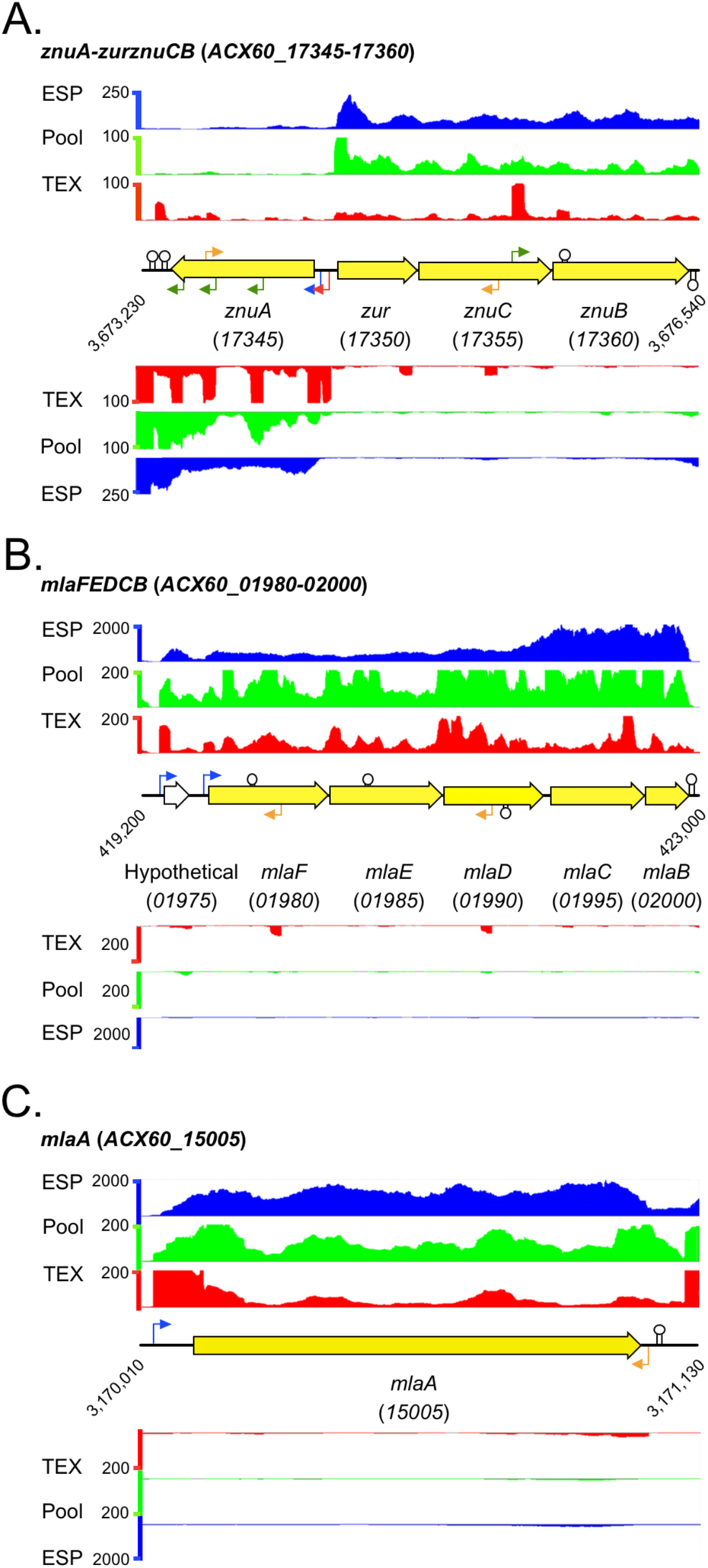
RNA-seq analysis of *znu* and *mla* virulence operon gene expression. Mapped, normalized RNA-seq reads illustrate the quantitative measure of expression. Right-angle arrows indicate TSS, which are colour-coded to correspond with the TSS categories described in figure 2. The lollipop structures represent predicted rho-independent transcriptional terminators.

#### The Membrane Lipid Asymmetry (Mla) Transport System

Several of the genes responsible for *A. baumannii* persistence in the mouse pneumonia model were identified as belonging to efflux pump systems (44). The authors highlighted the Mla ATP-binding cassette (ABC) transport system because this system was annotated as conferring a toluene tolerance phenotype (48), raising the question of why resistance to organic solvents may be important during infection of a mammalian host. However, recent studies have since uncovered a central role for Mla-mediated transport in maintaining the integrity of the outer membrane of Gram-negative bacteria (49, 50). Wang *et al*. reported a 15-fold decrease in *in vivo* persistence of mutants lacking *mlaA* (44), which correlates with roles for the Mla transport system in iron limitation and outer membrane vesicle formation as well as resistance to antibiotics such as doripenem and colistin (51–53).

Our transcriptomic data reveal that genes encoding the Mla ABC transporter components and periplasmic binding protein are co-transcribed in the *mlaFEDCB* operon (Fig. 5B). Similar to the *zurznuCB* operon, TSSpredator did not call a TSS upstream of this operon, but manual inspection of the TEX track revealed a single TSS located 29 nt upstream of *mlaF* at position 419,650. We noted greater transcript abundance of *mlaDCB* (most notably *mlaCB*) compared to *mlaFE* in ESP, an expression pattern that was also observed for an LPS-deficient mutant of *A. baumannii* strain 19606 (51). As there is no additional promoter within the operon, this suggests that the *mlaDCB* portion of the transcript may turnover more slowly or is protected from degradation resulting in elevated production of the periplasmic substrate binding domain MlaC. A rho-independent terminator was identified downstream of the operon and provides a potential 3’ border for the transcript; however, multiple rho-independent terminators were also identified within the operon, with two terminators present early in the *mlaF* and *mlaE* ORFs. Like *E. coli* and other Gram-negative bacteria, the *A. baumannii mlaA* homolog is found at a separate location on the chromosome (Fig. 5C) and we identified a single TSS for *mlaA*, located 74 nt upstream of the start codon. An additional antisense TSS was also identified, located opposite of the 3’ end of the *mlaA* ORF. Both the *mlaFEDCB* and *mlaA* loci were expressed during ESP, suggesting that this growth condition could be used for further investigation of the transport system.

#### Chloramphenicol resistance is regulated by transcription initiation

We discovered that *A. baumannii* 17978 cultured in M9 medium loses resistance to chloramphenicol (Fig. 6). All *A. baumannii* strains contain the chromosomally-encoded efflux pump CraA, for chloramphenicol resistance *Acinetobacter*. Efflux by CraA appears to be highly specific to chloramphenicol, conferring a minimum inhibitory concentration >256 μg mL^−1^ chloramphenicol (54). Inhibition of CraA in *A. baumannii* strain 19606 increases sensitivity by 32-fold while deletion of *craA* increases sensitivity by 128-fold, respectively (54). Expression of *craA* is highly consistent between strains of multi-drug resistant *A. baumannii* (55), but it is not known if this expression is constitutive in nature (56). In *A. baylyi*, a point mutation 12 bp upstream of the *craA* start codon caused increased stability of the *craA* mRNA resulting in higher *craA* expression and increased resistance to chloramphenicol (57).

**Figure 6.**
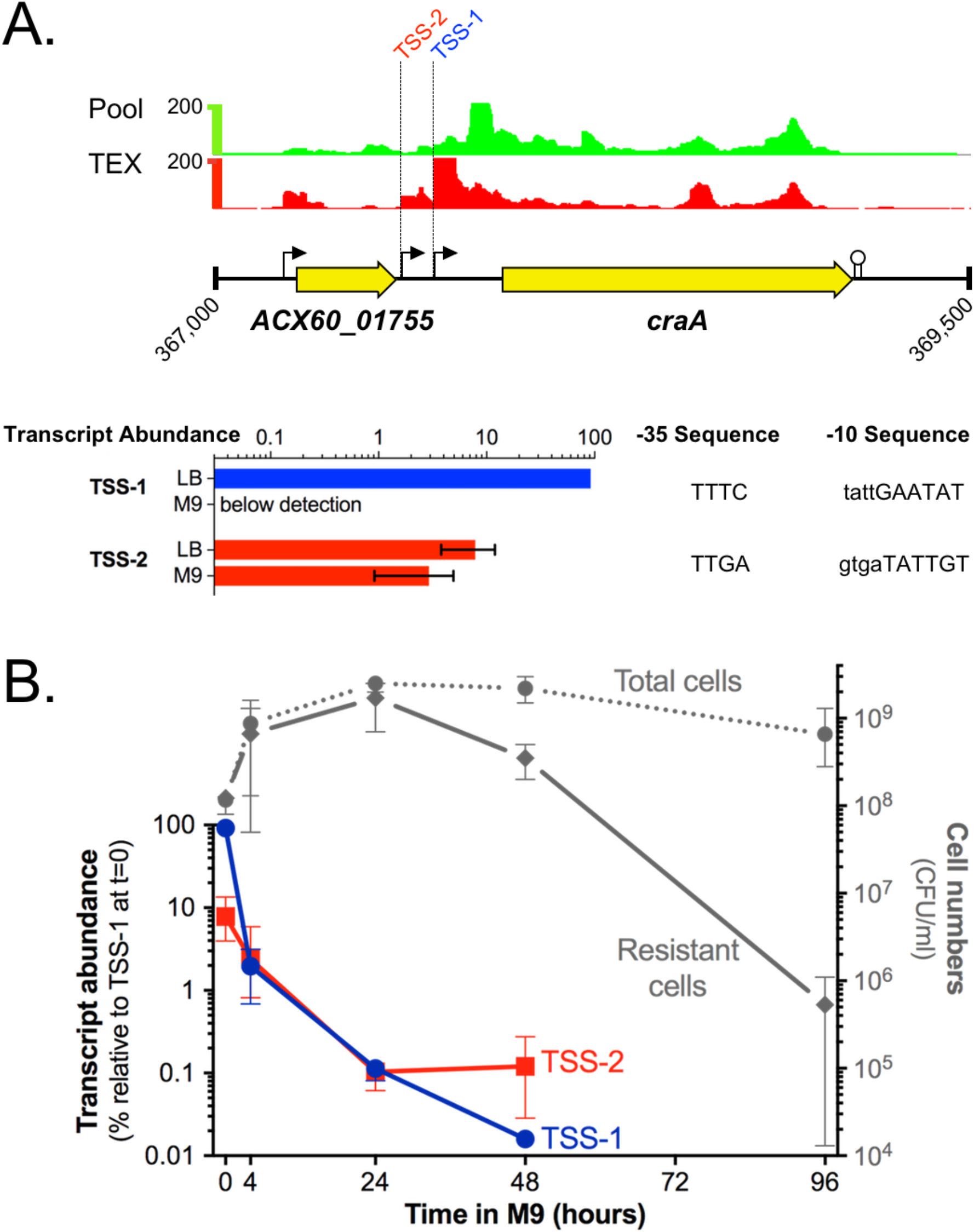
Elucidation of transcriptional control of chloramphenicol resistance. (**A**) Normalized, mapped sequence reads from dRNA-seq of the RNA pool (upper panel). Right-angle arrows indicate the two TSS identified in the promoter region of the chloramphenicol efflux pump gene, *craA*. The lollipop structure represents a predicted rho-independent transcriptional terminator. Transcript abundance was quantified for TSS-1 or TSS-2 and expressed relative to total transcript abundance in cells growing exponentially in LB medium, with predicted −10 and −35 RNAP binding sites. (B) Growth experiment of *A. baumannii* in M9 over 96 hours. At the indicated time points, *A. baumannii* cells were plated on LB agar (Total cells) or LB + chloramphenicol (Resistant cells). Transcript abundance of *craA* mRNA originating from TSS-1 or TSS-1+2 measured by qPCR at 0, 4, 24 and 48 hours.

We used dRNA-seq and quantitative PCR to examine the promoter architecture and regulation of *craA* in *A. baumannii* ATCC 17978, and tested whether loss of chloramphenicol resistance is coupled to altered expression of *craA* (Fig. 6A). dRNA-seq revealed two promoters at the *craA* gene, the gene-proximal TSS-1 promoter (position 367,680, Table S2) and the gene distal TSS-2 promoter (position 367,564, Table S2). To examine how *craA* expression is regulated in response to growth conditions, we used qPCR to differentiate transcripts originating at TSS-1 from TSS-2. Because TSS-1 is contained within TSS-2 transcripts, TSS-1 transcripts were calculated by subtracting TSS-2 transcripts from total *craA* transcripts. During steady-state rapid growth (exponential phase) in nutrient-rich L-broth, 92% of *craA* transcription originates at TSS-1 (Fig. 6A). Conversely, in the nutrient-poor medium M9 plus pyruvate, TSS-1 activity was undetectable and all transcription was driven by TSS-2. Next, we tested how the two promoters responded to nutrient downshift. Within four hours of transferring exponentially-growing cells from L-broth to M9 plus pyruvate, TSS-1 was down-regulated by 50-fold, whereas TSS-2 was down-regulated only 1.4-fold (Fig. 6B). Therefore, the promoter driving TSS-1 is highly responsive to nutritional quality of the growth medium. Over the following days, both promoters continued to decline in activity. TSS-1 has a poor match to the RpoD −35 consensus, suggesting that protein transcription factors control TSS-1 by recruiting RNAP to the promoter. The TSS-2 promoter contains an archetypical −35 sequence, suggesting that RNAP can bind this promoter without assistance from an activator protein, which could explain the consistent expression of TSS-2 in L-broth and M9 medium.

We hypothesized that down-regulation of *craA* in M9 medium would result in a chloramphenicol-sensitive phenotype akin to that of a *craA* mutant strain. Thus, we evaluated the proportion of resistant and sensitive *A. baumannii* cells in liquid cultures. During exponential growth in rich medium (L-broth), all cells were resistant to chloramphenicol (Fig. 6B). All cells remained resistant at four hours after transfer to M9 medium, but 24 hours after transfer to minimal media, 50% of cells were susceptible. Cell viability was high after 96 hours in M9 medium, but chloramphenicol resistance had declined by more than 3,000-fold (Fig. 6B). Taken together, these results suggest that antibiotic resistance is conditional, especially when cells are instantly deprived of amino acids upon transfer from Lennox medium to M9 plus pyruvate medium. Although chloramphenicol resistance is ubiquitous in *A. baumannii* isolates due to drug efflux by CraA, we have discovered that *craA* expression—and thus chloramphenicol resistance— is conditional on nutritional quality of the environment. This lack of endogenous activation or repression of drug resistance presents a potential target for the effective treatment of multi-drug resistant infections with an inexpensive and widely available antibiotic.

## Conclusion

As an emerging pathogen in the “omics” age, *A. baumannii* pathobiology is defined largely by high-throughput technologies that can guide molecular studies of pathogenicity mechanisms and antimicrobial resistance. We have conducted the first dRNA-seq analysis of the model strain *A. baumannii* ATCC 17978 to identify several thousand TSS at single nucleotide resolution, and generated a comprehensive map of the transcriptome that includes 110 sRNAs. Although 3731 TSS is the largest and most precise dataset yet for *Acinetobacter*, it is important to note that computational identification of TSS can overlook true sites. TSS are only called where a sharp increase in read depth in the TEX-treated sample exceeds the normalized read depth in the Pool sample. Overall, we identified an average of 5.1 sense strand TSS/10 kbp in *A. baumannii* ATCC 17978, which is similar but fewer than the 7.0 sense strand TSS/10 kbp identified in S. Typhimurium 4/74 (11). Because manual curation of S. Typhimurium 4/74 identified more TSS per bp, we suspect the Predator program used here did not identify all TSS. Thus, scientists using AcinetoCom are encouraged to examine the RNA-seq read depth at their gene(s) of interest. A sharp increase in read depth in the TEX track alone can be highly suggestive of a TSS. Thus, investigators are advised to scrutinize cases where there is high expression in the ESP tracks but no TSS is called in the TEX track.

A growing body of evidence suggests a positive relationship between conditions that slow bacterial growth and the expression of virulence factors. For notable pathogens such as *Legionella, Yersinia*, and *Salmonella*, key virulence traits are induced during the early stationary phase in laboratory culture (10, 58–61). Expression of virulence determinants in standard laboratory conditions has permitted the study of gene function and characterization of the regulatory networks that control virulence gene expression in other model pathogens (62). It was fortuitous to discover that *A. baumannii* expresses key pathogenicity genes in standard laboratory conditions, including the three loci identified in the mouse model of pneumonia (4, 44) highlighted in Figure 5. Similarly, simple laboratory conditions revealed an environmental dimension to the control of drug resistance in *A. baumannii*. Although diagnostic techniques that test for the presence of antibiotic resistance genes can predict resistance, our finding of chloramphenicol sensitivity in a species that is considered highly chloramphenicol resistant illustrates the importance of characterizing gene expression and gene regulatory mechanisms. Reduced chloramphenicol efflux due to rapid down-regulation of *craA* expression presents two intriguing leads for combating this ubiquitous resistance mechanism in *A. baumannii*. First, characterization of the regulatory mechanism and identification of the environmental cues that stimulate or repress *craA* expression will raise the potential to predict stages of infection and locations in the host that *A. baumannii* may be sensitive to chloramphenicol. Second, elucidation of the transcription factors controlling *craA* may reveal additional drug targets that could be targeted synergistically. This could enhance the effectiveness of a widely available and inexpensive antibiotic that is being used effectively in the treatment of other multi-drug resistant infections.

## Acknowledgements

This work was supported by funding from Saskatchewan Health Research Foundation (grant 2867) and Natural Sciences and Engineering Research Council of Canada (Discovery grant RGPIN-435784-2013) to ADSC. Saskatchewan Health Research Foundation (grant 3866) to KDM.

---

**Table 1. Growth Conditions Used to Induce the Global *A. baumannii* Transcriptome**

**Table S1. Oligonucleotides Used in this Study**

**Table S2. Transcriptional Start Sites of *A. baumannii* str. 17978-mff**

**Table S3. sRNAs of *A. baumannii* str. 17978-mff**

**Table S4. Expression data**

